# Loss of glutathione redox homeostasis impairs proteostasis by inhibiting autophagy-dependent protein degradation

**DOI:** 10.1101/309849

**Authors:** David Guerrero-Gómez, José Antonio Mora-Lorca, Beatriz Sáenz-Narciso, Francisco José Naranjo-Galindo, Fernando Muñoz-Lobato, Cristina Parrado-Fernández, Ángel Cedazo-Minguez, Christopher D. Link, Christian Neri, María Dolores Sequedo, Rafael P. Vázquez-Manrique, Elena Fernández-Suárez, Veit Goder, Roser Pané, Elisa Cabiscol, Peter Askjaer, Juan Cabello, Antonio Miranda-Vizuete

## Abstract

In the presence of aggregation-prone proteins, the cytosol and endoplasmic reticulum (ER) undergo a dramatic shift in their respective redox status, with the cytosol becoming more oxidized and the ER more reducing. However, whether and how changes in the cellular redox status may affect protein aggregation is unknown. Here, we show that *C. elegans* mutants lacking glutathione reductase *gsr-1* gene enhance the deleterious phenotypes of heterologous human as well as endogenous worm aggregation-prone proteins. These effects are phenocopied by the GSH depleting agent diethyl maleate. Additionally, *gsr-1* mutants abolish the nuclear translocation of HLH-30/TFEB transcription factor, a key inducer of autophagy, and strongly impair the degradation of the autophagy substrate p62/SQST-1::GFP, revealing glutathione reductase may have a role in the clearance of protein aggregates by autophagy. Blocking autophagy in *gsr-1* worms expressing aggregation-prone proteins results in strong synthetic developmental phenotypes and lethality, supporting the physiological importance of glutathione reductase in the regulation of misfolded protein clearance. Furthermore, impairing redox homeostasis in both yeast and mammalian cells induces toxicity phenotypes associated with protein aggregation. Together, our data reveal that glutathione redox homeostasis may be central to proteostasis maintenance through autophagy regulation.

## INTRODUCTION

The maintenance of a functional proteome (also known as proteostasis or protein homeostasis) requires the concerted intervention of different enzymatic and regulatory systems, termed as Proteostasis Network (PN). Thus, PN assists polypeptides to achieve and maintain their active conformation (chaperones and folding factors), provides an adequate cellular environment for their function (unfolded protein response (UPR), heat shock and oxidative responses) and efficiently disposes them when recognized as non-functional or are no longer required (proteasome and autophagy systems) ^1,2^. Given its pivotal role in cellular function, deregulation of PN dramatically impacts protein homeostasis and is an underlying cause of several human diseases, collectively known as proteinopathies, mainly characterized by the aberrant deposition of aggregated, misfolded proteins ^3^. These disorders include some of the most prevalent neurodegenerative diseases like Alzheimer’s, Parkinson’s or Huntington’s Diseases, among others ^4^. At the core of PN, the pathways involved in reduction, oxidation and isomerization of disulfide bonds are essential to maintain protein homeostasis as they prevent, correct or remove faulty bonds leading to non-functional protein folding.

Protein disulfide bonds are formed at cysteine residues, which can also undergo a variety of other post-translational modifications such as sulfenylation, sulfinylation, nitrosylation or persulfidation, some of which are irreversible ^5^. The maintenance of protein thiols in their reduced state in the cytosol and mitochondrial matrix is carried out mainly by two dedicated redox pathways: the thioredoxin and glutathione systems^6^. In contrast, in the lumen of the endoplasmic reticulum (the subcellular compartment where the folding of the proteins entering the secretory pathway takes place), the correct disulfide bond formation is performed by an oxidative folding pathway, largely composed of members of the protein disulfide isomerase (PDI) family ^7^. Intriguingly, no members of the thioredoxin or glutathione redox systems exist in the endoplasmic reticulum, and the identity of the enzymatic systems providing reducing equivalents to ER-resident PDIs has remained elusive for decades. Recently, the cytoplasmic thioredoxin system has been shown to shuttle electrons into the ER to reduce oxidized PDIs and ensure correct disulfide formation ^8^. In addition, an active import of cytoplasmic glutathione into ER lumen by specific transporters is key to maintain redox homeostasis in the ER ^9^, highlighting the importance of a coordinated redox network for maintaining proteostasis across all subcellular compartments.

Organisms as distant as the nematode *C. elegans* and humans contain a similar number of cysteine residues in their proteome, estimated in about 210,000 ^10^. Importantly, their respective thioredoxin and glutathione redox systems are highly conserved ^11^, supporting the use of *C. elegans* as a simple model relevant to addressing key questions in redox biology ^12,13^. In worms, the thioredoxin system is dispensable for redox homeostasis as null or strong loss of function mutants of the genes encoding the cytoplasmic and mitochondrial thioredoxins (*trx-1, trx-2* and *trx-3*) and thioredoxin reductases (*trxr-1* and *trxr-2*) are viable and superficially wild type ^14, 15, 16, 17^. In contrast, mice lacking any member of the cytosolic or mitochondrial thioredoxin system are embryonic lethal (reviewed in ^18^). However, when bypassing the embryonic requirement by employing conditional knock-outs, the thioredoxin system is also dispensable for viability in mice ^18^. Thus, in the absence of a functional thioredoxin system, the glutathione system may be responsible for the maintenance of redox homeostasis in both organisms ^16, 18,19^. This key role of glutathione is further emphasized by the strict requirement of dietary methionine as a GSH source (generated by the trans-sulfuration pathway) in the absence of functional thioredoxin and glutathione redox systems in mice ^19^ or the lethal phenotypes of mutations in the genes responsible for glutathione synthesis (*gcs-1* and *gss-1*) or recycling (*gsr-1)* in worms ^10, 20^. Furthermore, a study in *C. elegans* showed that GSH redox potential is highly sensitive to small changes in its oxidation status ^10^. This sensitivity would translate into rapid responses to even minor perturbations of the cytosolic redox environment and, as consequence, to a quick adjustment of the thiol-disulfide balance of the proteome. Together, these data strongly suggest the glutathione system is the primary enzymatic system responsible for the maintenance of redox homeostasis in metazoa.

Under non-stressed conditions, the cytosol of eukaryotic cells is maintained in a reduced state to favour the stabilization of free thiol groups while the ER environment is more oxidized in order to promote disulfide bond formation ^21^. Consistent with a tightly controlled interplay between proteostasis and redox homeostasis, proteotoxic stress generated by aggregating proteins in the cytoplasm of both *C. elegans* and mammalian cells strongly disturb redox homeostasis, causing a shift towards a more oxidizing condition in the cytosol and, conversely, towards a more reducing condition in the ER ^22^. However, despite the obvious influence of the redox environment in protein folding, whether disruption of redox homeostasis promotes protein aggregation is unknown. Here, we addressed this question by exploring the impact of a compromised glutathione redox environment in *C. elegans* models of aggregation-prone proteins. We have found that impairment of glutathione homeostasis causes a robust enhancement of phenotypes associated to protein aggregation as a consequence of autophagy disruption and we show this effect is evolutionary conserved in eukaryotes from yeast to mammalian cells.

## RESULTS

### *gsr-1* deficiency exacerbates the phenotypes induced by polyQ expression in *C. elegans* muscle cells

Although proteotoxic stress may trigger loss of redox homeostasis ^22^, how impaired redox homeostasis impacts on protein aggregation is unknown. To address this question, we focused on the role of the *C. elegans gsr-1* gene, encoding glutathione reductase, for two reasons: Firstly, the glutathione redox system is essential in worms ^10, 20^ whereas the thioredoxin system is dispensable ^14, 15, 16, 17^. Secondly, maternal *gsr-1* mRNA and/or protein contribution from heterozygous progenitors (balanced with the qC1 balancer ^23^) allows to work with genetically homozygous *gsr-1* animals, designated as *gsr-1(m+,z-)*, because these animals reach adulthood with no discernible phenotype under normal growth conditions ^20^, in contrast with *gss-1* and *gcs-1* mutants that are unable to synthesize glutathione and in which maternal contribution only allows growth of the homozygous progeny till L1-L2 larval stages ^10^.

On this premise, we first downregulated *gsr-1* expression by RNAi feeding in worms that constitutively express in muscle cells heterologous aggregation-prone proteins associated to human neurodegenerative diseases such as *α*-synuclein::YFP (Parkinson) ^24^, Q40::YFP (polyglutamine diseases like Huntington) ^25^ and human Aβ (Alzheimer) ^26^. Despite to the fact that A*β, α*-synuclein and polyQ stretches lack cysteine residues in their sequence, *gsr-1* downregulation increased the number of *α*-synuclein::YFP and Q40::YFP aggregates while it enhanced the paralysis of Aβ worms (a readout of Aβ aggregation) (data now shown). This suggests that the phenotypes caused by *gsr-1* downregulation are likely not mediated by direct disulphide bond formation on these proteins. Unexpectedly, worms expressing Q40::YFP in muscle cells displayed a progressive larval arrest phenotype when feeding on *gsr-1* RNAi bacteria, an effect that did not occur in transgenic animals expressing the control Q35::YFP fusion protein ^25^ (**Supplementary Figure 1a,b**) or in the Aβ and *α*-synuclein::YFP models (data not shown). Consistent with the RNAi data, *gsr-1(m+,z-)* animals had a significantly increased number of muscle Q40::YFP aggregates at the L4 (day 3) and young adulthood (day 4) stages (**Figure 1a**), which inversely correlated with the muscular performance measured by thrashing activity in liquid medium (**Supplementary Figure 1c**). No effect was found on the aggregation or thrashing phenotypes of *rmIs132 [Punc-54::Q35::yfp]; gsr-1(m+,z-)* control worms (**Figure 1a** and data not shown). Transgenic restoration of GSR-1 expression in *rmIs133 [Punc-54::Q40::yfp]; gsr-1(m+,z-)* worms decreased the number of Q40::YFP aggregates, demonstrating that the increased aggregation phenotype is specific to *gsr-1* deficiency and not to a closely linked unknown mutation (**Supplementary Figure 1d**). Furthermore, pharmacological inhibition of GSH synthesis (by buthionine sulfoximine, BSO), GSSG reduction (by 1,3-bis(2-chloroethyl)-1-nitrosourea, BCNU) and GSH depletion (by diethyl maleate, DEM) phenocopied the developmental delay phenotype caused by *gsr-1* downregulation in worms expressing Q40::YFP fusion protein (**Supplementary Figure 1e,f and Figure 1b**). Remarkably, DEM had a much more profound effect than BSO or BCNU as *rmIs133 [Punc-54::Q40::yfp]* hermaphrodites exposed to 2.5 mM DEM rapidly became sick and laid very few progeny which invariably arrested at L1-L2 larval stage (**Figure 1b**). Together, these data demonstrate that reduced GSR-1 and GSH levels compromise muscle function and survival of worms expressing Q40::YFP in muscle cells.

**Figure 1.**
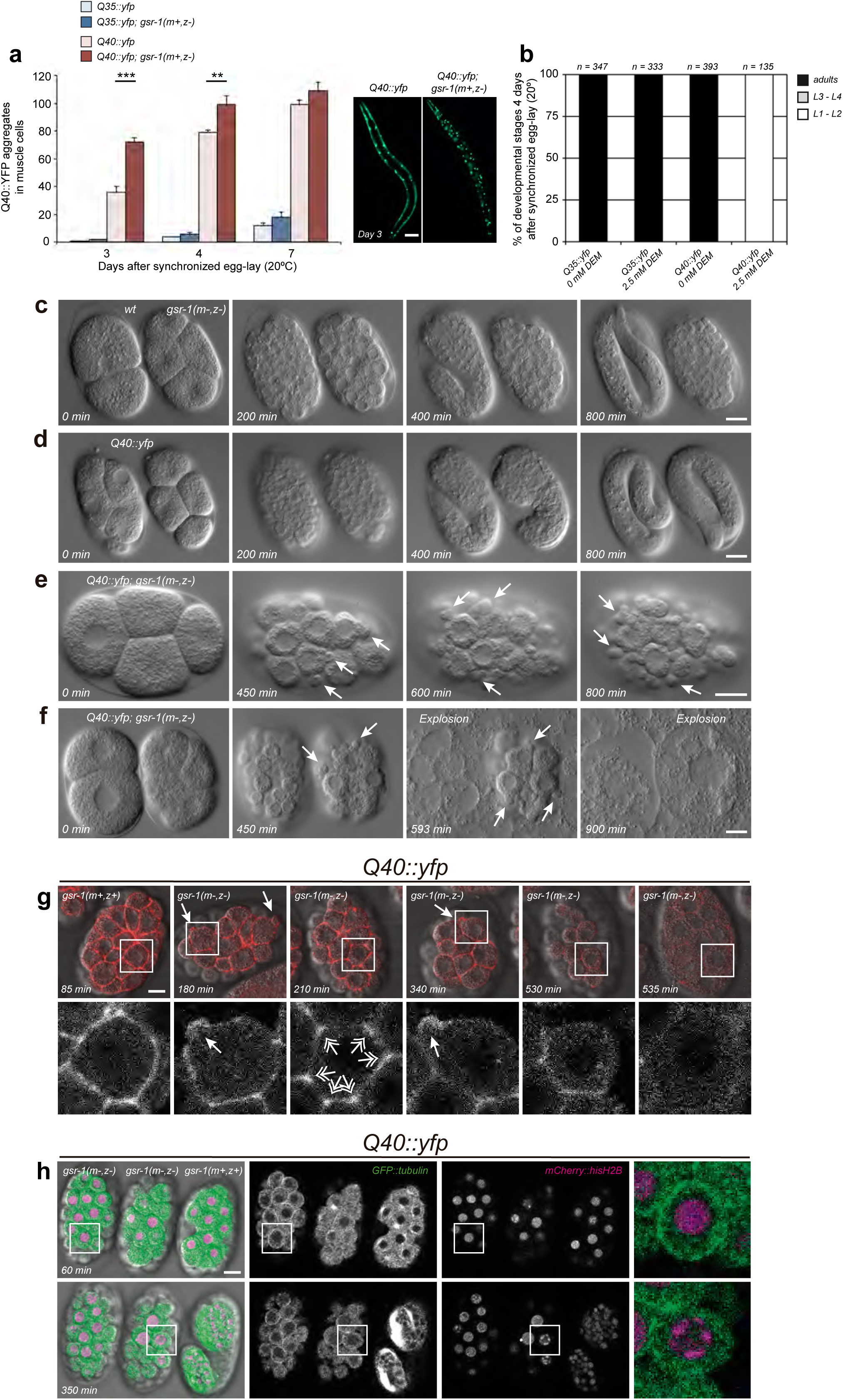
Phenotypes of the *gsr-1* mutation and DEM treatment on *C. elegans* expressing Q40::YFP protein in muscle cells. (**a**) The *gsr-1(m+,z-)* mutation increases the number of Q40::YFP, but no Q35::YFP, aggregates in worm muscle cells. Data are the mean ± S.E.M. of 3 independent experiments (n=10 animals per strain and assay). ** *p*<0.01; \*\*\**p*<0.001 by unpaired, two-tailed Student’s *t*-test. Images show one representative example of muscle Q40::YFP aggregates for each genotype. Scale bar 100 μm. (**b**) 2.5 mM DEM treatment causes a fully penetrant L1-L2 larval arrest on worms expressing a Q40::YFP fusion protein in muscle cells while no effect is found in *Q35::yfp* animals. Data are from 3 independent experiments (n=total number of animals assayed). (**c-f**) Differential interference contrast still images of developing *wild type* and *gsr-1(m-,z-)* embryos (**c**), *Q40::yfp* embryos (**d**) and *Q40::yfp; gsr-1(m-,z-)* embryos (**e,f**). Arrows indicate transient blebs. Embryo explosion in (**f**) is denoted by the sudden disappearance of embryonic cells. The still images in **c-f** are taken from Movies 1 to 4, respectively. (**g**) Control *gsr-1(m+,z+)* and *Q40::yfp; gsr-1(m-,z-)* embryos expressing LifeAct::mCherry were mounted and recorded together. Bottom row shows higher magnification images of LifeAct::mCherry only. Closed arrows point to examples of membrane blebs whereas double-headed arrows indicate F-actin accumulation at cell vertices. Last frame corresponds to the moment when the eggshell suddenly fills completely and the embryo explodes. The still images are taken from Movie 5. (**h**) Control *Q40::yfp; gsr-1(m+,z+)* and *Q40::yfp; gsr-1(m-,z-)* embryos expressing GFP::tubulin and mCherry::hisH2B (green and magenta in merge, respectively) were mounted and recorded together. Right column shows higher magnification images of boxed area. Strong accumulation of microtubules at the plasma membrane is observed at 60 min (top row). Condensation of chromatin at the nuclear periphery occurs at 350 min. Additional examples are provided in Movie 6. All movie recordings were carried out at 25°C in H_2_O except in (**h**) that was performed in M9 buffer at 25°C. Time is indicated from start of recording. Scale bar 10 μm.

### *gsr-1(m-,z-)* embryos expressing Q40::YFP in muscle cells undergo dramatic cell blebbing and exploding phenotypes

We previously reported that *gsr-1(m-,z-)* embryos, lacking maternal contribution, arrest at the pregastrula/gastrula stage with the embryonic cells superficially normal and no apparent signs of necrosis or apoptosis (**Figure 1c and Movie 1**) ^20^. In turn, *rmIs133 [Punc-54::Q40::yfp]* embryos develop normally until hatching, similar to wild type embryos (**Figure 1d and Movie 2**) ^25^. Surprisingly we identified a robust synthetic phenotype in *rmIs133 [Punc-54::Q40::yfp]; gsr-1(m-,z-)* embryos as most of these embryos undergo dramatic cell blebbing and/or sudden catastrophic embryo explosion phenotypes when maintained in water at 25°C (**Figure 1e,f, Movies 3-4 and Table 1**). These phenotypes also happened, although to a lesser extent, in *gsr-1(m-,z-)* embryos expressing a Q35::YFP fusion protein from the *rmIs132 [Punc-54::Q35::yfp]* transgene (**Table 1**). Importantly, when these embryos are maintained in an isotonic buffer, they do not explode but maintain the blebbing phenotype, suggesting the eggshell integrity, which preserves embryonic osmotic homeostasis, is compromised in *gsr-1(m-,z-)* embryos expressing Q35::YFP or Q40::YFP proteins (**Table 1**).

**Table 1:**
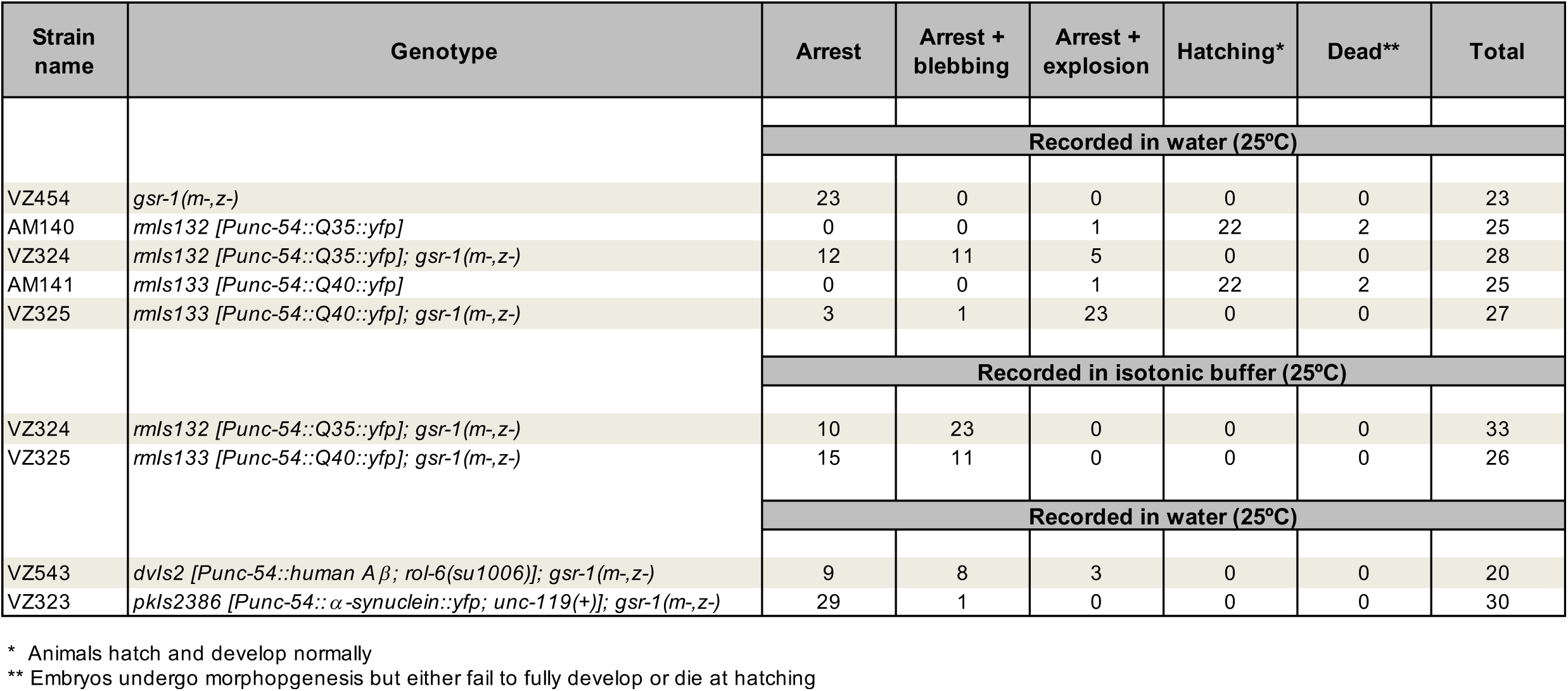
Phenotypes of embryos expressing different aggregation proteins in muscle cells.

Under normal conditions, the plasma membrane of eukaryotic cells is tightly bound to an underlying actomyosin cortex that contributes to cell shape maintenance ^27^. However, when the actomyosin cortex integrity is disrupted, actin-devoid evaginations (blebs) are generated by internal hydrostatic pressure ^28^. To test whether impairment of the actomyosin cortex function might underlay the blebbing phenotype of the *rmIs133 [Punc-54::Q40::yfp]*; *gsr-1(m-,z-)* embryos, we examined the dynamics of actin fluorescence markers. The distribution of actin filaments in *rmIs133; gsr-1(m-,z-)* embryos did not markedly differ from that of *rmIs133; gsr-1(m+,z+)* controls and actin was also detected at the sites of blebs (**Figure 1g and Movie 5)**. As microtubule network destabilization can also result in cell blebbing ^29,^,^30^, we next asked whether tubulin dynamics was altered in *rmIs133; gsr-1(m-,z-)* embryos. Using tubulin fluorescence markers, we identified an aberrant distribution of the tubulin filaments in the periphery of the *rmIs133; gsr-1(m-,z-)* embryonic cells that is not observed in control embryos (**Figure 1h and Movie 6**). Similar results were obtained with tubulin immunostaining (**Supplementary Figure 2**). These data suggest that, in a GSH compromised environment, polyQ proteins modify the dynamics of the microtubule network, impinging on plasma membrane integrity and function.

**Figure 2.**
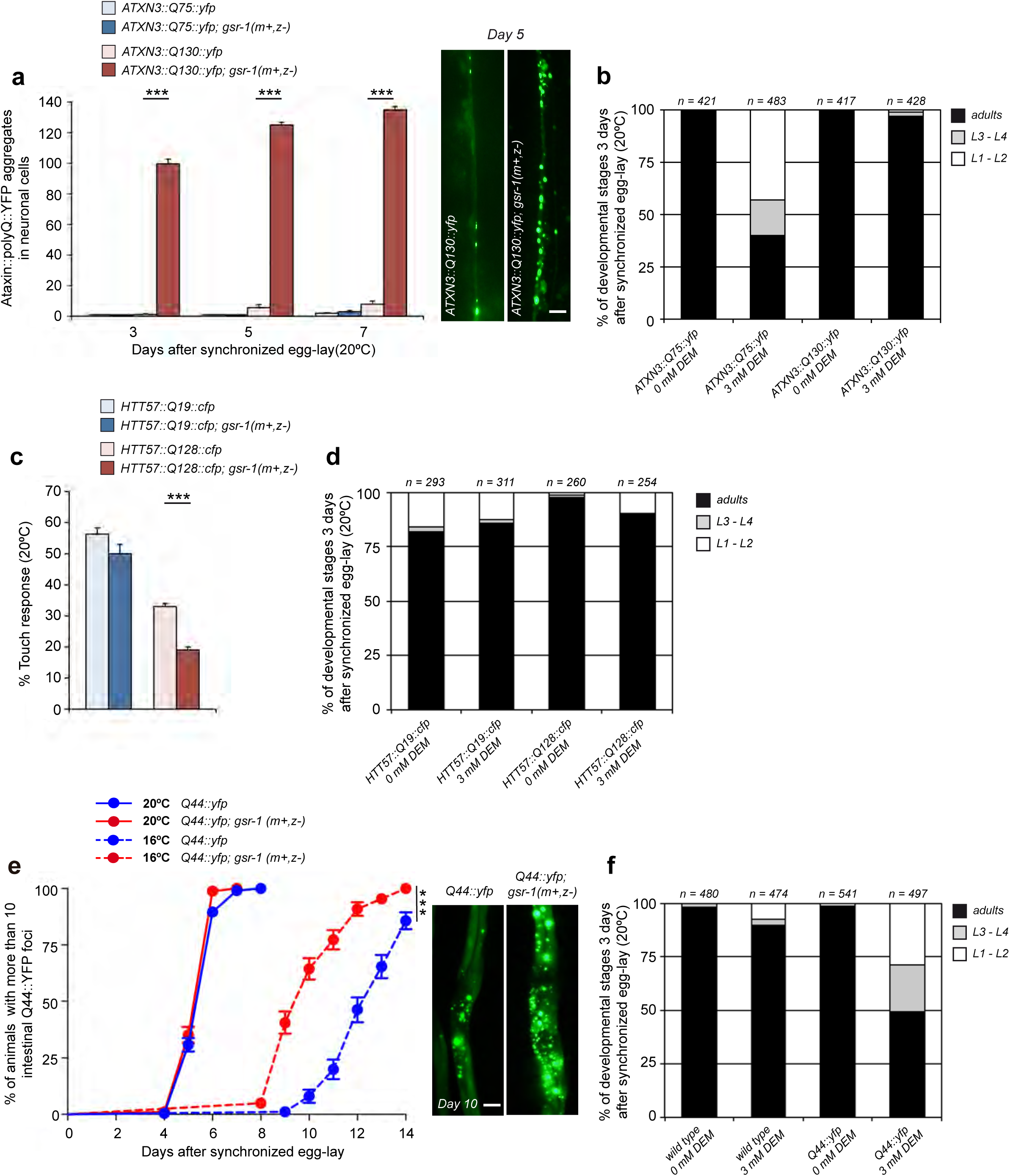
Phenotypes of the *gsr-1* mutation and DEM treatment on *C. elegans* expressing polyQ proteins in neurons and intestinal cells. (**a**) The *gsr-1(m+,z-)* mutation increases the number of ATXN3::Q130::YFP aggregates in the worm ventral nerve cord while no effect is found in control worms expressing ATXN3::Q75::YFP fusion protein. Data are the mean ± S.E.M. of 3 independent experiments (n=12 animals per strain and assay). \*\*\**p*<0.001 by unpaired, two-tailed Student’s *t*-test. Images show one representative example of ventral nerve cord ATXN3::Q130::YFP aggregates for each genotype. Scale bar 20 μm. (**b**) 3 mM DEM treatment delays the larval development of worms expressing pan-neuronal ATXN3::Q75::YFP but not ATXN3::Q130::YFP fusion proteins. Data are from 3 independent experiments (n=total number of animals assayed). (**c**) Percentage of touch-responsiveness of worms expressing HTT57::Q19::CFP and HTT57::Q128::CFP fusion proteins in mechanosensory neurons of *wt* versus *gsr-1(m+,z-)* worms. Data are the mean ± S.D. of 2 independent experiments (n*≥*100 animals per strain and assay). \*\*\**p*<0.001 by One-way ANOVA with Tukey’s multiple comparison test. (**d**) 3 mM DEM treatment does not affect the larval development of worms expressing HTT57::Q19::CFP and HTT57::Q128::CFP fusion proteins in mechanosensory neurons. Data are from 3 independent experiments (n=total number of animals assayed). (**e**) Q44::YFP aggregates onset of appearance in *wt* versus *gsr-1(m+,z-)* worms as function of time. Data are the mean ± S.E.M of 3 independent experiments (n=50 animals per strain and assay). \*\*\**p*<0.001 by Log-rank (Mantel-Cox) test. Images show one representative example of intestinal Q44::YFP aggregates for each genotype. Scale bar 50 μm. (**f**) 3 mM DEM treatment delays the larval development of worms expressing a Q44::YFP fusion protein in intestinal cells. Data are from 3 independent experiments (n=total number of animals assayed).

### *gsr-1* mutants show enhanced polyQ proteins aggregation in neurons and intestinal cells

To determine whether the protective effect of GSR-1 in polyQ worms may be restricted to muscle cells or extended to other tissues, we analyzed *gsr-1(m+,z-)* animals expressing polyQ proteins in neuronal and intestinal cells. In addition, to pharmacologically validate the data obtained with *gsr-1(m+,z-)* mutants in these models, we opted for depleting the pool of reduced glutathione by DEM treatment as it produced the more penetrant phenotype on *rmIs133 [Punc-54::Q40::yfp]* worms (**Figure 1b**). First, we used a *C. elegans* model of Machado-Joseph Disease pathogenesis in which mutant human Ataxin-3 (ATXN3) fused to YFP is expressed in a pan-neuronal fashion ^31^. Similar to muscular polyQ models, we found that the *gsr-1(m+,z-)* mutation caused a robust increase of ventral nerve cord fluorescent foci in worms expressing mutant ATXN3::Q130::YFP but not in ATXN3::Q75::YFP worms that served as non-aggregating controls (**Figure 2a**). However, upon exposure of animals to DEM, we observed an inverse correlation with toxicity as the ATXN3::Q75::YFP control worms suffered a significant developmental delay while, in contrast, ATXN3::Q130::YFP worms developed at the same rate compared to untreated controls (**Figure 2b**). These results suggest that a protective mechanism might operate in neurons when a certain aggregation threshold is surpassed.

We further confirmed the deleterious effect of the *gsr-1* mutation on neuronal polyQ toxicity using worms that express in the six touch receptor neurons the first 57 amino acids of the human huntingtin protein followed by normal (Q19) or expanded (Q128) polyglutamines and that is fused to CFP ^32^. While the *gsr-1* mutation did not modify light touch response of worms expressing the HTT57::Q19::CFP fusion protein, we observed a significant decrease of the touch response in HTT57::Q128::CFP nematodes (**Figure 2c**). In contrast to the ataxin model, we did not identify any differential growth response of HTT57::Q19::CFP and HTT57::Q28::CFP worms to DEM treatment (**Figure 2d**), most likely due to the dispensability of touch neurons for development.

Finally, we also assayed the impact of the *gsr-1* mutation in worms expressing a Q44::YFP fusion protein in intestinal cells ^33^. At 20°C, the appearance onset of intestinal fluorescent aggregates was similar in both wild type and *gsr-1(m+,z-)* backgrounds (**Figure 2e**). However, at 16°C, the formation of intestinal Q44::YFP aggregates occurred earlier and these aggregates were bigger and more abundant in *gsr-1(m+,z-)* worms compared to wild type controls (**Figure 2e**). In consonance, animals expressing Q44::YFP proteins were highly sensitive to DEM (**Figure 2f**). Collectively, these data indicate that *gsr-1* and GSH deficiency enhance polyQ proteins aggregation regardless of the cell type or tissue of expression.

### *gsr-1* deficiency is deleterious in *C. elegans* expressing human Aβ and *α*-synuclein proteins

In addition to the effect on polyQ proteins aggregation, the previously mentioned enhanced paralysis of worms expressing human Aβ or increased *α*-synuclein::YFP aggregation in animals fed with *gsr-1* RNAi pointed to a general effect of *gsr-1* in proteostasis. We then characterized the effect of the *gsr-1* mutation in these two additional models of protein aggregation. Worms expressing the *dvIs2 [Punc-54::Aβ]* transgene produce human Aβ protein in muscle cells where it aggregates and causes age-dependent progressive paralysis at 20°C but not (or very mildly) at the 16°C permissive temperature ^26^. As shown in **Figure 3a**, the *gsr-1(m+,z-)* mutation induced robust paralysis in *dvIs2 [Punc-54::Aβ]* worms at the permissive temperature. This synthetic interaction was phenocopied pharmacologically with DEM, causing a complete L1-L2 larval arrest phenotype of *dvIs2* worms (**Figure 3b**). Moreover, *dvIs2 [Punc-54::Aβ]*; *gsr-1(m-,z-)* embryos also showed blebbing and explosion phenotypes (**Table 1**), similar to *gsr-1(m-,z-)* embryos expressing Q40::YFP under the control of the muscle specific *unc-54* promoter (**Figure 1e,f**). Interestingly, an even more striking phenotype was found when combining the *gsr-1(m+,z-)* mutation with a different Aβ transgene, *dvIs14 [Punc-54::Aβ]* ^34^. In this case, *dvIs14*; *gsr-1(m+,z-)* worms at the permissive temperature displayed a much faster paralysis as compared to that of *dvIs2; gsr-1(m+,z-)* counterparts (**Figure 3c**). In addition, *dvIs14*; *gsr-1(m+,z-)* animals showed a fully penetrant egg-laying phenotype, most likely reflecting the lack of contractility of the uterine and vulva muscles which are responsible for egg extrusion (**Figure 3d**). The genetic interaction between Aβ protein and the *gsr-1* mutation was also found in worms expressing human Aβ in the nervous system from the *ganIs2 [Punc-119::Aβ]* transgene ^35^. Thus, the *gsr-1(m+,z-)* mutation causes a strong developmental delay in a *ganIs2* genetic background (**Figure 3e**) which correlated with a higher sensitivity to DEM of *ganIs2* nematodes (**Figure 3f**).

**Figure 3.**
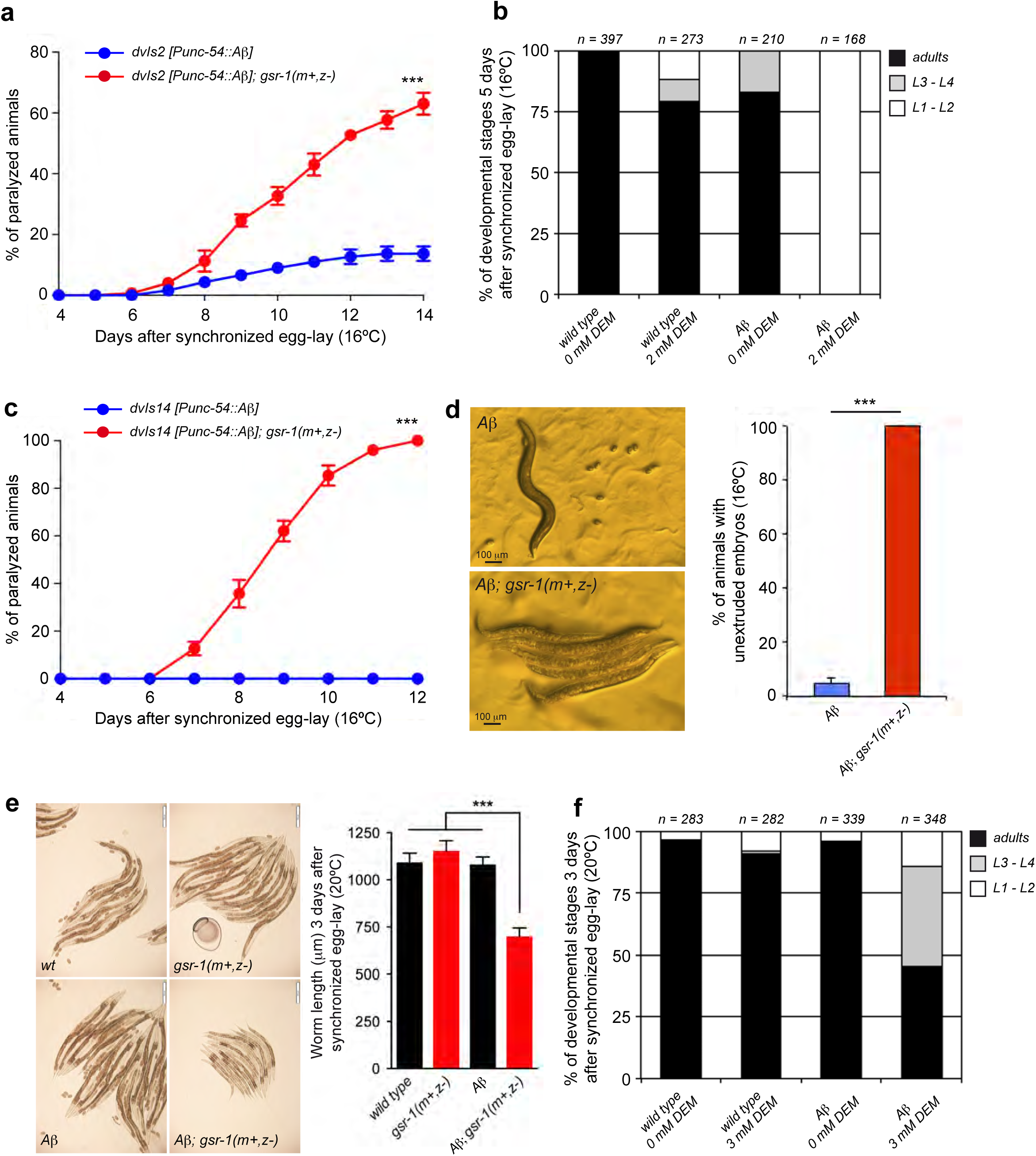
Phenotypes of the *gsr-1* mutation and DEM treatment on *C. elegans* expressing the human *β*-amyloid peptide. (**a**) The *gsr-1(m+,z-)* mutation increases the paralysis of worms expressing the human *β*-amyloid peptide in muscle cells from the *dvIs2* integrated array. Data are the mean ± S.E.M. of 3 independent experiments (n=25 animals per strain and assay). \*\*\**p*<0.001 by Log-rank (Mantel-Cox) test. (**b**) 2 mM DEM treatment causes a fully penetrant L1-L2 larval arrest on worms expressing the human *β*-amyloid peptide in muscle cells. Data are from 3 independent experiments (n=total number of animals assayed). (**c**) When expressing the human *β*-amyloid peptide in muscle cells from the *dvIs14* integrated array, the *gsr-1(m+,z-)* mutation increases the paralysis of worms much faster than that observed in *dvIs2* animals. Data are the mean ± S.E.M. of 3 independent experiments (n=25 animals per strain and assay). \*\*\**p*<0.001 by Log-rank (Mantel-Cox) test. (**d**) The *gsr-1(m+,z-)* mutation provokes a fully penetrant unextruded embryo phenotype in worms expressing the human *β*-amyloid peptide in muscle cells from the *dvIs14* array. Data are the mean ± S.E.M. of 3 independent experiments (n=50 animals per strain and assay). \*\*\**p*<0.001 by unpaired, two-tailed Student’s *t*-test. Images show one representative example of *dvIs14* worms in a *wt* or *gsr-1(m+,z-)* background. Scale bar 100 μm. (**e**) The *gsr-1(m+,z-)* mutation delays the development of worms expressing the human *β*-amyloid peptide in the nervous system. Data are the mean ± S.D. of 40 worms per genotype. \*\*\**p*<0.001 by unpaired, two-tailed Student’s *t*-test. Images show one representative example for each genotype. Scale bar 200 μm. (**f**) 3 mM DEM treatment delays the larval development of worms expressing the human *β*-amyloid peptide in neurons. Data are from 3 independent experiments (n=total number of animals assayed).

Likewise, the aggregation in muscle cells of the *α*-synuclein::YFP fusion protein expressed from the *pkIs2386 [Punc-54::α-syn::yfp]* transgene ^24^ was strongly enhanced in *gsr-1(m+,z-)* mutants (**Figure 4a**) although, in contrast to animals expressing Q40::YFP or Aβ in muscle cells, DEM treatment did not cause any developmental phenotype in muscle *α*-synuclein::YFP animals (**Figure 4b**). Similarly, in *baIn11 [Pdat-1::α-synuclein; Pdat-1::gfp]* animals expressing *α*-synuclein in the six worm head dopaminergic neurons ^36^, the *gsr-1(m+,z-)* mutation significantly increased neurodegeneration in an age-dependent manner (**Figure 4c**) whereas no phenotype was observed when these animals are treated with DEM (**Figure 4d**). The lack of any deleterious effect of DEM treatment in worms expressing *α*-synuclein::YFP in muscle cells may be due to the very mild effect on motility which has been reported for these worms ^24^ in comparison to human Aβ or Q40::YFP expressing animals ^25, 26^. Similarly, the dispensability of dopaminergic signalling for worm survival ^37^ may explain the absence of deleterious phenotypes of worms expressing *α*-synuclein in dopaminergic neurons when exposed to DEM.

**Figure 4.**
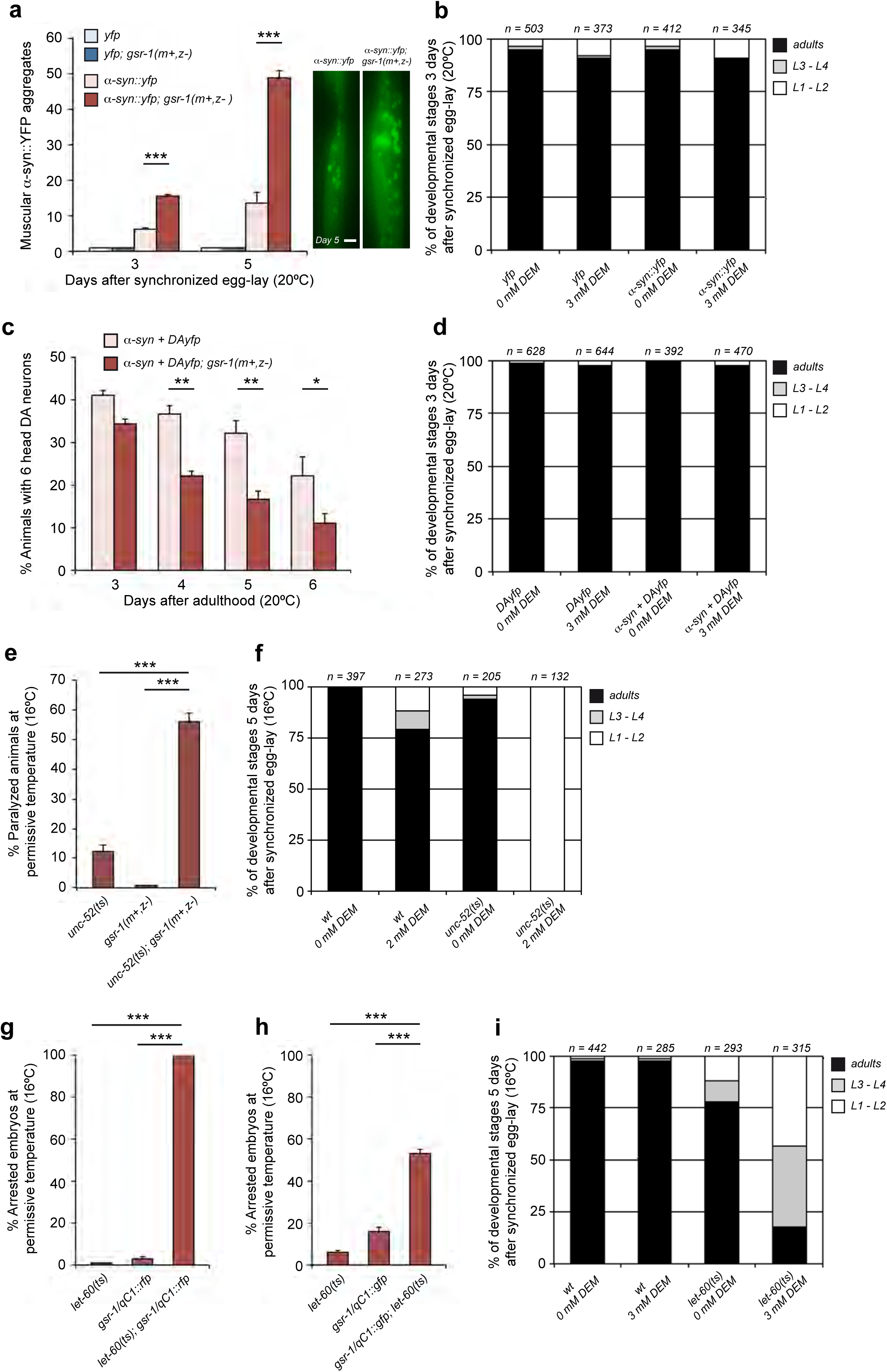
Phenotypes of the *gsr-1* mutation and DEM treatment on *C. elegans* expressing human *α*-synuclein and endogenous metastable proteins. (**a**) The *gsr-1(m+,z-)* mutation increases the number of *α*-SYN::YFP aggregates in the muscle cells. Aggregate quantification was restricted to aggregates located between the two pharyngeal bulbs. Data are the mean ± S.E.M. of 3 independent experiments (n=12 animals per strain and assay). \*\*\**p*<0.001 by unpaired, two-tailed Student’s *t*-test. Images show one representative example of muscle *α*-SYN::YFP aggregates for each genotype. Scale bar 10 μm. (**b**) 3 mM DEM treatment does not affect the development of worms expressing the *α*-SYN::YFP fusion protein in muscle cells. Data are from 3 independent experiments (n=total number of animals assayed). (**c**) The *gsr-1(m+,z-)* mutation accelerates the age-dependent neurodegeneration of dopaminergic neurons expressing human *α*-synuclein. Data are the mean ± S.E.M. of 3 independent experiments (n=30 animals per strain and assay). \**p*<0.05, \*\**p*<0.01 by unpaired, two-tailed Student’s *t*-test. (**d**) 3 mM DEM treatment does not affect the development of worms expressing human *α*-synuclein in dopaminergic neurons. Data are from 3 independent experiments (n=total number of animals assayed). (**e**) The *gsr-1(m+,z-)* mutation increases the paralysis phenotype of *unc-52(e669su250)* worms at permissive temperature. Data are the mean ± S.E.M. of 3 independent experiments (n≥70 animals per strain and assay, scored at day 7 after egg-lay). \*\*\**p*<0.001 by unpaired, two-tailed Student’s *t*-test. (**f**) 2 mM DEM treatment causes a fully penetrant L1-L2 larval arrest on *unc-52(e669su250)* worms at permissive temperature. Data are from 3 independent experiments (n=total number of animals assayed). (**g,h**) The *gsr-1(m+,z-)* mutation increases the embryonic arrest phenotype of *let-60(ga89)* worms at permissive temperature. (**g**) *let-60(ga89); gsr-1/qC1::rfp* worms produced only arrested embryos while (**h**) *let-60(ga89); gsr-1/qC1::gfp* worms only produced both balanced viable progeny and arrested embryos but not unbalanced *gsr-1(m+,z-)* progeny. Data are the mean ± S.E.M. of 3 independent experiments (n≥100 embryos/animals per strain and assay). \*\*\**p*<0.001 by unpaired, two-tailed Student’s *t*-test. (**i**) 3 mM DEM treatment delays the larval development of *let-60(ga89)* worms at permissive temperature. Data are from 3 independent experiments (n=total number of animals assayed).

### *gsr-1* deficiency worsens the phenotypes of *C. elegans* expressing endogenous metastable proteins

All models so far described in this work employ heterologous aggregation-prone proteins (Aβ, a-synuclein::GFP or polyQ::GFP) not naturally found in *C. elegans*. To rule out that the effect of the *gsr-1(m+,z-)* mutation in these models might be a consequence of expressing foreign proteins in worm cells, we monitored mutant phenotypes in animals bearing thermosensitive alleles of endogenous genes, which encode metastable proteins that are highly sensitive to changes in the protein folding environment. To this end, we used *unc-52(e669su250)* and *let-60(ga89)* mutants that display a highly penetrant paralysis or embryonic arrest phenotype, respectively, at the non-permissive temperature of 25°C but are roughly wild type at the permissive temperature of 16°C ^38^. Consistent with the results above, the double mutant *unc-52(e669su250); gsr-1(m+,z-)* displayed a robust paralysis phenotype at 16°C (**Figure 4e**). This synthetic interaction was phenocopied by treatment of *unc-52(e669su250)* mutants with 2mM DEM, causing a fully penetrant early larval arrest (**Figure 4f**). Likewise, a strong synthetic interaction was found between *let-60(ga89)* and *gsr-1* mutants as shown by the impossibility to generate viable progeny from *let-60(ga89); gsr-1/qC1::rfp* balanced animals at permissive temperature (**Figure 4g**). To rule out a possible deleterious effect of the *qC1::rfp* balancer, we stabilized the *gsr-1(tm3574)* mutation with a different version of the *qC1* balancer tagged with GFP (see Supplementary Table 1 for full genomic information of both balancers) that resulted in 50% viable progeny (**Figure 4h**). Remarkably, none of this viable progeny was *gsr-1(m+,z-),* but all *gsr-1(m+,z+)*, thus confirming the synthetic lethal phenotype of *gsr-1(m+,z-)* and *let-60(ga89)* mutants which was further supported by the larval arrest phenotype of *let-60(ga89)* animals in the presence of 3mM DEM (**Figure 4i**). Together, these data suggest that glutathione reductase and the maintenance of a proper GSH redox homeostasis may protect against protein aggregation.

### Autophagy function is impaired in *gsr-1* mutants

Proteostasis is maintained by a delicate equilibrium of different molecular pathways. Some of these pathways are at the core of the proteome maintenance such as protein synthesis, folding, trafficking or degradation while others act as modifiers of proteostasis such as unfolded protein response, nutritional and metabolic status, stress situations, genetic and epigenetic susceptibility or physiological processes like aging ^2^. To ascertain which of these pathways might underlay the protective effect of GSR-1 and GSH in protein aggregation, we first tested the effect of *gsr-1* deficiency on different reporters of proteostasis regulators (including insulin, proteasome, UPR-ER, heat-shock, mitophagy or autophagy pathways), under basal and stress conditions.

As shown in **Supplementary Figure 3**, the *gsr-1(m+,z-)* mutation only caused a mild downregulation of the UPR-ER reporter HSP-4::GFP under both conditions (**Supplementary Figure 3c**) and an induction of the oxidative stress reporter SOD-3::GFP under basal, but not stressed conditions (**Supplementary Figure 3b**). As SOD-3::GFP induction was not accompanied by an increase in DAF-16::GFP nuclear translocation (**Supplementary Figure 3a**), the effect of the *gsr-1* mutation on *sod-3* transcription is more likely due to a DAF-16 independent response to oxidative stress rather than a role in proteostasis.

In contrast to all reporters described in **Supplementary Figure 3**, we identified a clear effect of the *gsr-1* mutation in worms carrying the *sqIs17* transgene, that expresses the HLH-30::GFP autophagy marker ^39^. HLH-30 is the worm orthologue of the mammalian TFEB transcription factor and, under normal growth conditions, these transcription factors are inactive and diffusely distributed in the cytoplasm. However, upon stress conditions such as starvation and heat-shock, HLH-30/TFEB translocate into the nucleus, triggering the transcription of genes required for autophagy induction and lysosomal biogenesis ^39, 40, 41, 42^. We found that the cytosolic subcellular localization of HLH-30::GFP was not modified in a *gsr-1(m+,z-)* mutant background in well-fed animals (**Figure 5a**). In turn, one hour starvation or 30°C heat-shock was enough to induce a strong nuclear translocation of HLH-30::GFP, a phenomenon severely hampered in *gsr-1(m+,z-)* worms subjected to starvation, but not heat-shock (**Figure 5a and Supplementary Figure 4a**). These results suggest that *gsr-1* deficiency may negatively impact on proteostasis by impairing autophagy-dependent protein degradation, at least upon specific stimuli. To test for this possibility, we used worms that carry the *bpIs151* transgene and express SQST-1::GFP, a reporter protein that is typically degraded by autophagy in non-stressed conditions and that accumulates in the form of fluorescent foci, mainly in embryonic and larval intestinal cells, when autophagy is compromised ^43^. Consistently, we found no SQST-1::GFP foci in *bpIs151; gsr-1(m+,z-)* embryos and L1 larvae under normal growth conditions (**Figure 5b,c**). However, 24 hours starvation or a mild impairment of autophagy by RNAi downregulation of *lgg-1*/LC3 (required for autophagosome biogenesis) increased the number of SQST-1::GFP foci in *gsr-1(m+,z-)* embryos and L1 larvae, respectively, as compared to wild type controls (**Figure 5b,c and Supplementary Figure 4b**). These results identify GSR-1 as a novel positive regulator of autophagy.

**Figure 5.**
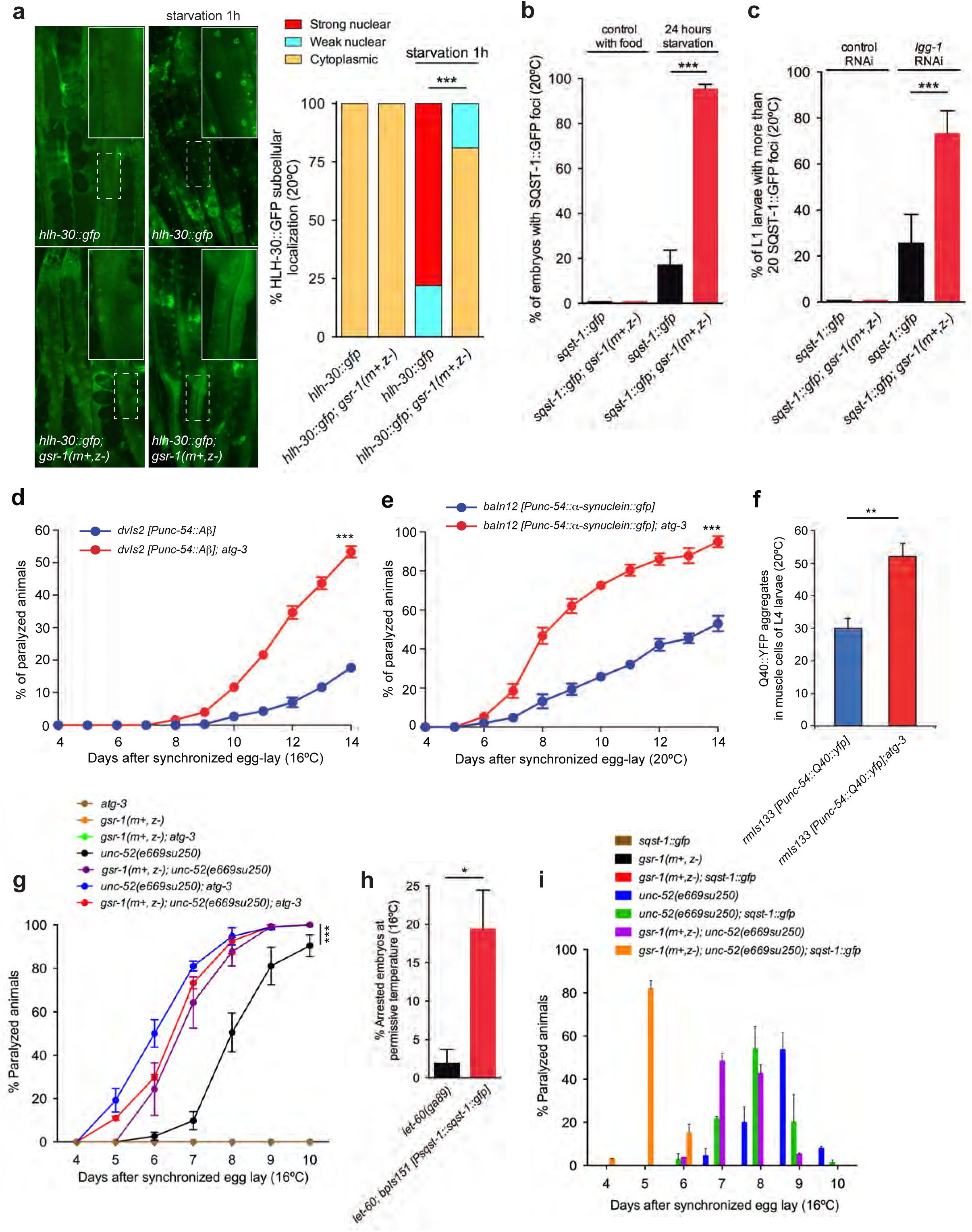
The *gsr-1* mutation enhances the phenotypes of *C. elegans* proteostasis models by preventing autophagy function. **(a)** The *gsr-1(m+,z-)* mutation does not modify the diffuse cytoplasmic localization of the HLH-30::GFP reporter under well-fed conditions but strongly inhibits its nuclear translocation after one hour starvation. Data are the mean of 3 independent experiments (n=50 animals per strain and assay). \*\*\**p*<0.001 by Chi^2^ test. (**b-c**) The *gsr-1(m+,z-)* mutation increases the appearance of SQST-1::GFP foci in (**b**) embryos generated by parents that have been starved for 24 hours and (**c**) in L1 larvae fed with *lgg-1* RNAi. Data are the mean of 2 independent experiments (n=50 animals per strain and assay). \*\*\**p*<0.001 by unpaired, two-tailed Student’s *t*-test. (**d,e**) The *atg-3(bp412)* mutation increases the paralysis of (**d**) worms expressing the human *β*-amyloid peptide in muscle cells from the *dvIs2* integrated array and (**e**) of worms expressing *α*-synuclein::GFP in muscle cells from the *baIn12* integrated array. Data are the mean ± S.E.M. of 3 independent experiments (n=25 animals per strain and assay). \*\*\**p*<0.001 by Log-rank (Mantel-Cox) test. (**f**) The *atg-3(bp412)* mutation increases the number of Q40::YFP aggregates in worm muscle cells. Data are the mean ± S.E.M. of 3 independent experiments (n=10 animals per strain and assay). ** *p*<0.01 by unpaired, two-tailed Student’s *t*-test. (**g**) The *atg-3(bp412)* mutation increases the paralysis of *unc-52(e669su250)* worms, expressing metastable UNC-52 protein in muscle cells. Data are the mean ± S.E.M. of 3 independent experiments (n *≥*50 animals per strain and assay). \*\*\**p*<0.001 by Log-rank (Mantel-Cox) test of all strains carrying the *unc-52(e669su250)* allele versus the *unc-52* single mutant control. (**h**) Expression of the SQST-1::GFP autophagy substrate increases the embryonic arrest phenotype of *let-60(ga89)* worms. Data are the mean ± S.E.M. of 3 independent experiments (n≥100 embryos/animals per strain and assay). \*\*\**p*<0.001 by unpaired, two-tailed Student’s *t*-test. (**i**) Expression of the SQST-1::GFP autophagy substrate increases the paralysis onset of *unc-52(e669su250)* worms and this effect is further enhanced in a *gsr-1(m+,z-)* background. Data are the mean ± S.E.M. of 3 independent experiments (n*≥*50 animals per strain and assay). *p*<0.001 by Log-rank (Mantel-Cox) test of all strains carrying the *unc-52(e669su250)* allele versus the *unc-52* single mutant control.

Next, we reasoned that if *gsr-1* mutants enhance the phenotypes of worm models of proteostasis by impairing autophagy, then genetically blocking autophagy in these proteostasis models would similarly result in an enhancement of their deleterious phenotypes. To our knowledge, no reports have so far addressed the implication of autophagy in the phenotypes of *unc-52* or *let-60* thermosensitive mutants while a role for autophagy on the phenotypes of worms expressing human A*β, α*-synuclein and polyQ proteins has only been studied by RNAi downregulation ^42, 44, 45, 46^. To provide more compelling evidence of the implication of autophagy in these proteostasis models, we employed *atg-3(bp412)* mutants, in which autophagy is blocked at the autophagosome formation step ^43^. We corroborated the RNAi results in the muscle A*β, α*-synuclein and polyQ worm models and found an enhancement of their respective paralysis or aggregation phenotypes in an *atg-3* background (**Figure 5d-f**). Interestingly, we also found a significant enhancement of the paralysis in *unc-52; atg-3* double mutants compared to *unc-52* controls at permissive temperature (**Figure 5g**), a result that was phenocopied using *lgg-1* RNAi (**Supplementary Figure 4c**). We also attempted to evaluate the effect of the *atg-3(bp412)* mutation in the embryonic arrest phenotype of *let-60(ga89)* mutants but we failed to isolate a double mutant *atg-3; let-60*, which possibly indicates a lethal interaction. In support of this possibility, *let-60(ga89)* mutants expressing the autophagy substrate SQST-1::GFP display a ten-fold increase in the embryonic arrest phenotype at the permissive temperature (**Figure 5h**). Together, these data reveal that the endogenous, aggregation-prone metastable UNC-52 and LET-60 proteins are likely degraded by autophagy and reinforce the implication of autophagy in dismissing heterologous A*β, α*-synuclein and polyQ proteins in worm models of neurodegenerative diseases.

Glutathione depletion has been shown to increase protein carbonylation and aggregation ^47^. We hypothesized that the deleterious effect of the *gsr-1* mutation on proteostasis could be explained by an increase of the load of proteins to be degraded by autophagy, ultimately resulting in a severe impairment of the autophagy machinery. To test this hypothesis, we aimed to generate *atg-3(bp412)* and *gsr-1(m+,z-)* double mutants in the different protein-aggregation models and we faced strong synthetic interactions. For instance, we were unable to generate viable *atg-3; gsr-1(m+,z-)* animals expressing human Aβ protein in muscle cells (either from *dvIs2* or *dvIs14* transgenes). In consonance, although viable, *atg-3; gsr-1(m+,z-)* worms expressing *α*-SYN::GFP in muscle cells were sick and displayed a high percentage of internal hatching, which precluded proper quantification of paralysis (data not shown). Likewise, *atg-3; gsr-1(m+,z-)* animals expressing Q40::YFP in muscle cells were viable but had increased embryonic arrest phenotype (**Supplementary Figure 4d**). Only the triple *atg-3; gsr-1(m+,z-); unc-52* failed to show an increased paralysis phenotype compared to the two control strains *atg-3; unc-52* and *gsr-1(m+,z-); unc-52* (**Figure 5g**). However, increasing the autophagy load by expressing the SQST-1::GFP autophagy substrate in *unc-52; gsr-1(m+,z-)* worms resulted in a much earlier paralysis onset compared to *unc-52; gsr-1(m+,z-)* or *unc-52; bpIs151* double mutant controls (**Figure 5i**). Furthermore, a triple *atg-3; gsr-1/qC1::rfp; bpIs151* produced very few *gsr-1(m+,z-)* progeny which were very sick and arrested early during development (data not shown). Similarly, we were also unable to isolate triples *let-60; atg-3; bpIs151* or *let-60; gsr-1(m+,z-); bpIs151*, reinforcing the idea that metastable LET-60 is also degraded by autophagy. Together, the fact that *gsr-1(m+,z-)* mutants strongly enhance the deleterious phenotypes caused by aggregation-prone proteins in worms with compromised autophagy function (i.e. *atg-3* mutations or by expression SQST-1::GFP protein) pinpoints a key role of GSR-1 in maintaining a healthy proteome.

### Glutathione reductase mutation and glutathione depletion increase protein aggregation in yeast and mammalian cell cultures

To determine whether the protective role of the glutathione system on protein aggregation and associated phenotypes might be restricted to *C. elegans* or instead is a more universal, evolutionarily conserved mechanism, we turned to a *Saccharomyces cerevisiae* model of polyQ toxicity^48^. Importantly, the mechanism by which glutathione reductase regulates polyQ aggregation appears to be maintained as yeast cells lacking glutathione reductase (carrying a *Δglr1* deletion) and expressing the aggregation-prone Q103-GFP fusion protein displayed a clear aggregation pattern, not detected in the control strain expressing a non-aggregating Q25-GFP fusion protein (**Figure 6a**). Noteworthy, the aggregation phenotype of Q103-GFP expressing cells was much more evident when yeast have reached the diauxic phase (**Figure 6a**), when cells shift from a glucose fermentative to an ethanol respiration mitochondrial oxidative metabolism. Interestingly, protein aggregation of Q103-GFP was increased in *Δglr1* cells compared to a WT background only at diauxic phase, when oxidative stress generated by the metabolic shift occurred. In addition, *Q103-gfp* yeast had a significant growth delay in a *Δglr1* background and were more sensitive to DEM treatment than control Q25-GFP expressing cells (**Figure 6b-c**), reminiscent of the larval developmental arrest phenotype described for Q40::YFP worms with downregulated *gsr-1* levels (**Supplemental Figure 1a,b**) or exposed to DEM (**Figure 1b**). To test whether autophagy is also involved in maintaining yeast proteostasis when redox homeostasis is challenged, we generated a double *Δglr1Δatg8* mutant in *Q25-gfp* and *Q103-gfp* backgrounds. *atg8* encodes an ubiquitin-like protein which is essential for autophagosome formation ^49^. As shown in **Figure 6d**, single *Δglr1* or *Δatg8* mutants grew similar to wild type controls independently of expressing Q25-GFP or Q103-GFP proteins. However, a mild synthetic growth defect was detected in the double *Δglr1Δatg8* in both *Q25-gfp* and *Q103-gfp* backgrounds (**Figure 6d)** and this synthetic interaction was further enhanced when cells were exposed to 2.5 mM DEM **(Figure 6e)**. These data suggest that impairment of glutathione redox homeostasis negatively impacts on proteostasis in yeast cells expressing aggregation-prone proteins, involving autophagy as a mechanism to dismiss the increased load of protein aggregation.

**Figure 6.**
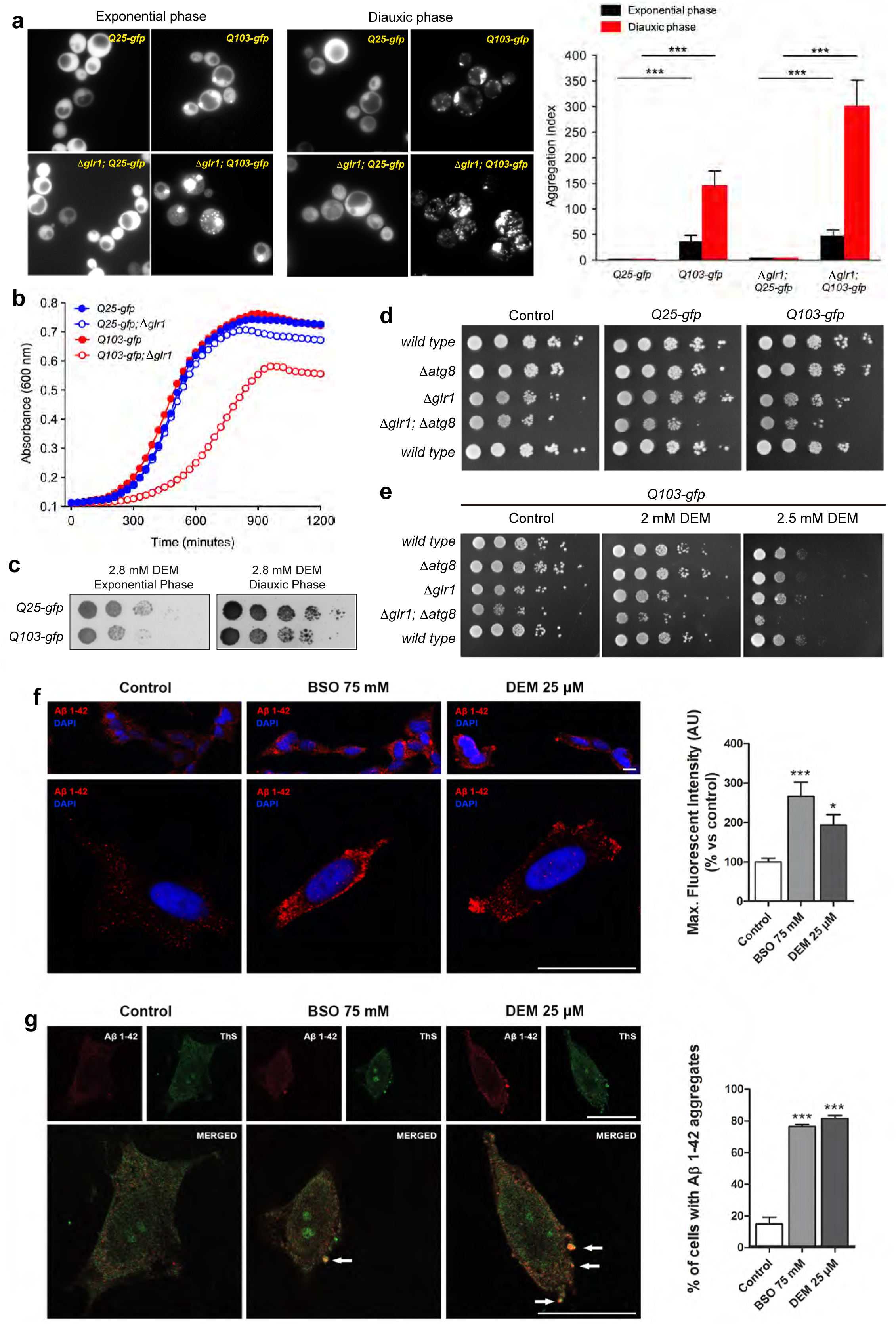
The protective role of the glutathione system in proteostasis maintenance is conserved from yeast to mammals. WT and *Δglr1* yeast strains expressing either 25Q-gfp or 103Q-gfp were grown in SC medium. (**a**). PolyQ-gfp fusion proteins were visualized by fluorescence microscopy and quantified (aggregation index) in cultures grown to exponential or diauxic phase as described in Materials and Methods. The *Δglr1* mutation increases the aggregation of 103Q-gfp but not 25Q-gfp. Data are the mean ± S.E.M. of 3 independent experiments with 4 biological replicates each. \*\**p*<0.01 by unpaired, two-tailed Student’s *t*-test. (**b**) Growth curves were measured in a microplate spectrophotometer for 20 h and only the *Δglr1;* 103Q-gfp strain shows growth delay. The data are the averages of two independent experiments with three biological replicates each. (**c**) Viability of Q25-gfp and Q103-gfp yeast measured by plating serial dilutions (1:10) of cells grown to exponential or diauxic phase in SC plates without or with DEM (2.8 mM). (**d,e**) Effect of *Δglr1* and/or *Δatg8* mutations on the viability of Q25-gfp and Q103-gfp yeast under (**d**) non-stressed conditions or (**e**) treated with DEM. Viability was measured by plating serial dilutions (1:10) of cells grown to diauxic phase prior to plating on SC plates. (**f**) Confocal microscopy images of SH-SY5Y cells incubated for 18 h in the presence of BSO (75 mM) and DEM (25 μM). A clear increase of intracellular Aβ1-42 staining is found in BSO and DEM treated cells. Aβ1-42 is shown in red and DAPI staining for nuclei is shown in blue. Data are the mean ± S.E.M. of 3 independent experiments (n=45 images with at least 5 cells/image per assay and treatment). \**p*<0.01, \*\*\**p*<0.001 by One way ANOVA followed by Newman-Keuls’ test. **(g)** The Aβ1-42 aggregates in cells treated with BSO and DEM (in red), colocalize with amyloid fibrils labeled by Thioflavin S staining (in green). Data are the mean ± S.E.M. of 3 independent experiments (n*≥*50 cells per assay and treatment). \*\*\**p*<0.001 by One way ANOVA followed by Newman-Keuls’ test.

Next, we addressed the protective effect of GSH homeostasis in a mammalian system using the SH-SY5Y neuroblastoma cell line stably transfected with the human APP protein, which drives intracellular accumulation of human Aβ ^50^. Interestingly, when exposing these cells to sublethal doses of BSO and DEM (**Supplementary Figure 5**), we found that both treatments promoted an increase of Aβ labelling in the cytoplasm of these cells (**Figure 6f**). Moreover, the number of cells with Aβ foci that colocalize with thioflavin, which labels amyloid fibrils and aggregates, was strongly induced upon BSO and DEM treatments (**Figure 6g**). Given that SH-SY5Y cells overexpressing APP have enhanced autophagy ^50^, the increased Aβ aggregation observed in BSO and DEM treated cells is consistent with an impairment of autophagy.

Collectively, these data confirm that the deleterious effect of GSH and glutathione reductase deficiency on protein aggregation may be conserved from lower eukaryotes to mammals, impinging on the autophagy machinery.

## Discussion

The maintenance of a functional proteome is essential for organismal survival and, among the molecular pathways that modulate proteostasis, redox regulation arises as a main regulatory mechanism allowing proteins to attain their native conformation and functionality. It has recently been proposed that the glutathione couple GSH/GSSG, rather than operating as a redox buffer would instead act as a surveillance relay to detect changes of the intracellular redox environment to tightly control the thiol-disulfide balance of the cellular proteome ^10^. In this study, we present a set of experiments that support this premise of the glutathione system being essential to preserve a healthy proteome, showing that genetic or pharmacological disruption of glutathione redox homeostasis enhances protein aggregation by a mechanism that impinges on the efficacy of autophagy.

### Loss of glutathione redox homeostasis enhances the toxicity of protein aggregation

Pharmacological inhibition of glutathione synthesis and recycling as well as glutathione depletion phenocopies the effect of the *gsr-1* mutation in worms expressing aggregation-prone proteins. Surprisingly, treatment with DEM in *C. elegans* expressing Aβ, Q40::YFP and metastable UNC-52 proteins in muscle cells caused a fully penetrant lethality phenotype. DEM is an alkylating agent that has been traditionally used to deplete GSH but a recent study has shown that DEM depletes not only GSH but also protein thiols^51^. Hence, it is plausible that DEM causes an overload of oxidatively compromised proteins, possibly deemed for autophagic degradation, in addition to its direct effect on the GSH/GSSG ratio. This double impact may underlie the enhanced phenotypes observed in the worm proteostasis models exposed to DEM as compared to those carrying the *gsr-1* mutation. Although it has been proposed that cytotoxicity can be genetically uncoupled from aggregation ^52^, we found a clear correlation of increased aggregation and DEM toxicity when aggregation-prone proteins are expressed in muscle cells. However, this correlation was not so strong for proteins aggregating in neurons. We indeed found one case in which cytotoxicity inversely correlates with aggregation, namely in worms that express the human ATXN3::Q130::YFP protein in neurons and that are highly resistant to DEM treatment despite undergoing extensive aggregation, contrasting with worms that express non-aggregating ATXN3::Q75::YFP protein which are very sensitive to DEM (**Figure 2a,b**). The resistance of ATXN3::Q130::YFP worms to the effects of DEM could be explained by an increase of exophers production, an extrusion mechanism mediated by very large extracellular vesicles containing damaged material such as aggregated proteins and that is enhanced when mitochondria are compromised ^53^, as is the case of *gsr-1* mutants ^20^. Plasma membrane blebbing can also be observed in body wall muscle cells of transgenic worms expressing human Aβ, and these blebs contain Aβ aggregates (CD Link, unpublished data). Therefore, body wall muscle cells might also generate exophers under specific conditions and tissue-specific penetrance of the *gsr-1* mutation or sensitivity to DEM treatment on exophers formation may account for the differential sensitivity of neuron versus muscle worm proteostasis models.

An unexpected phenotype identified in our experiments is the cell blebbing and explosion of *gsr-1(m-,z-)* embryos that carry the *rmIs133 [Punc-54::Q40::YFP]* or *dvIs2 [Punc-54::human Aβ]* transgenes. To our knowledge, such striking phenotypes have not been previously described in *C. elegans* embryos. Cell blebs arise from transient disruption of the actomyosin cortex underneath the plasma membrane ^28^ and we were surprised to find that actin network distribution did not differ obviously from that of non-blebbing controls. In turn, we found that microtubules were aberrantly distributed in the periphery of the *gsr-1(m-,z-)*; *rmIs133 [Punc-54::Q40::YFP]* embryonic cells. A recent report has identified the worm calponin CHDP-1 protein as promoter of cell protrusions in *C. elegans*. CHDP-1 is located at the cell cortex but, surprisingly, does not bind actin ^54^. Instead CHDP-1 associates with the Rho small GTPase family member Rac1/CED-10 which, among other functions, mediates microtubule dynamics and lamellipodial protrusions ^55, 56, 57^. Interestingly, we have found that *ced-10(t1875)* null mutant embryos also undergo cell blebbing, although these blebs are not as large and prominent as those observed in *gsr-1(m-,z-); rmIs133* embryos (**Movie 7**). Furthermore, *ced-10* mutants (both reduction and gain of function) increase *α*-synuclein::GFP and Q40::YFP aggregates in worm muscle cells and enhance *α*-synuclein dependent dopaminergic neurodegeneration (^58^ and A. Miranda-Vizuete, unpublished data), suggesting that CED-10 could mediate the protective effect of GSR-1 in these models. How could GSR-1 control CED-10 function? One possibility is via glutathionylation (a post-translational modification whereby a glutathione moiety oxidizes a specific cysteine residue ^59^), as mammalian Rac1 is glutathionylated at Cys18, a residue conserved in worm CED-10 protein, resulting in increased lamellipodia formation ^60^. Given that glutathionylation is carried out by glutaredoxins, a class of small redox proteins that are reduced by GSH ^61^, it will be very interesting to explore whether glutaredoxins mediate the protective effect of GSR-1 in proteostasis maintenance.

### Role of cytoplasmic versus mitochondrial glutathione reductase in proteostasis

Like the majority of eukaryotes, *C. elegans* uses the flavoenzyme glutathione reductase GSR-1 to recycle reduced glutathione from its oxidized form. In worms, *gsr-1* is an essential gene that encodes two different GSR-1 isoforms, one located in the cytosol and the other in mitochondria. While GSR-1 activity in the cytoplasm is absolutely required for embryonic development, consistent with glutathione synthesis occurring in this compartment, mitochondrial GSR-1 is dispensable under non-stressed conditions (^20^ and A. Miranda-Vizuete unpublished data). Surprisingly, *gsr-1(m+,z-)* animals exhibit mitochondria associated phenotypes such as mitochondrial fragmentation, induction of mitochondrial UPR and lowered mitochondrial DNA content ^20^. All the proteostasis models used in our study generate aggregated proteins in the cytosol, suggesting that the absence of GSR-1 activity in this compartment is responsible for the synthetic phenotypes observed in *gsr-1(m+,z-)* mutants. However, we cannot rule out a possible role of the mitochondrial GSR-1 isoform as mitochondria have been shown to import and degrade aggregated proteins originally produced in the cytoplasm ^62,63^. Because *gsr-1(m+,z-)* mutants display phenotypes associated to mitochondrial dysfunction, which might interfere with the import and degradation of the aggregated cytoplasmic proteins into this organelle, the implication of mitochondrial GSR-1 in proteostasis needs to be evaluated separately from the cytoplasmic isoform. Moreover, yeast Q103-GFP aggregates are more prominent in the diauxic phase, in which cells shift from a cytosolic fermentative to a mitochondrial respiratory metabolism, further supporting the possibility that mitochondrial glutathione reductase may have a role in protein aggregation.

### Glutathione-dependent regulation of autophagy

Autophagy is an evolutionarily conserved degradation system of defective proteins and subcellular structures, and an early event in autophagy induction is the nuclear translocation and transactivation of the basic Helix-Loop-Helix transcription factor HLH-30 ^39, 38^. Oxidative stress is one of the various conditions that activate autophagy (reviewed in ^64,,65^) and several components of the autophagy machinery are redox regulated by ROS-mediated mechanisms in different organisms ^66,67,68^. Using a HyPer fluorescent biosensor ^69^ we found that *gsr-1(m+,z-)* worms have wild type levels of H_2_O_2_ (A. Miranda-Vizuete, unpublished results), suggesting that they are not under oxidative stress and consistent with the lack of HLH-30::GFP nuclear translocation or increased SQST-1:GFP foci in these animals. Similarly, the *gsr-1* mutation does not interfere with HLH-30 nuclear translocation upon heat-shock, another ROS-producing treatment that induces autophagy ^42^ ^70^. In turn, *gsr-1(m+,z-)* mutants prevent HLH-30 nuclear translocation and degradation of the autophagy substrate p62/SQST-1 upon starvation, a condition that induces autophagy mainly through inhibition of the mTOR pathway ^71^. Together, these data suggest that the regulation of HLH-30 subcellular localization by GSR-1 is stimulus-specific, likely mediated by mechanisms that do not involve ROS signaling. Strikingly, *C. elegans* HLH-30 and mammalian TFEB proteins possess only one cysteine residue, located within (*C. elegans*) or in close proximity (mammalian) to their respective bHLH domains (**Supplementary Figure 6**). Similar to CED-10 regulation discussed above, glutathionylation of this only cysteine residue of HLH-30 could be a possible mechanism by which GSR-1 regulates HLH-30 nuclear translocation and transactivation, as it has been described for other transcription factors ^61, 72, 73^.

We previously showed that *gsr-1* mutants induce SKN-1 regulated genes of the glutathione pathway like *gcs-1* and *gst-4* ^20^. Moreover, GSR-1 levels are strongly upregulated upon starvation by a mechanism dependent on the SKN-1 transcription factor ^74^. Although SKN-1 may regulate mitophagy, a form of autophagy that specifically degrades defective mitochondria ^75^, it is not clear whether it can also regulate autophagy and protein aggregation in worms. Nonetheless, the Nrf2 transcription factor, which is the SKN-1 mammalian orthologue, is strongly associated to autophagy and proteotoxicity ^76, 77^, thus supporting the possibility that SKN-1 is a key player in the molecular mechanisms underlying GSR-1 function in these scenarios. Indeed, TFEB and Nrf2 transcription factors may cooperate to stimulate autophagy in mammalian cells^78^, making likely this cooperation may also happen in *C. elegans* for increasing GSR-1 levels upon starvation.

Additional evidence of GSR-1 impinging on autophagy comes from the fact that *gsr-1(m+,z-)* mutants further enhance the deleterious phenotypes caused by aggregation-prone proteins in worms with compromised autophagy function either by *atg-3* mutations or by overexpression of the SQST-1::GFP protein, which is specifically degraded by autophagy. This is nicely illustrated in *unc-52; gsr-1(m+,z-); sqst-1::gfp* worms, as they develop a much earlier onset of paralysis than the respective controls, probably by prioritizing SQST-1::GFP degradation and thus increasing the amount of aggregating UNC-52 that accelerates paralysis. Altogether, the synthetic phenotypes described in this study can be explained by a synergistic overload of misfolded proteins that cannot be degraded by autophagy, which ultimately surpasses a threshold over which cell function is severely affected eventually causing organismal death.

In summary, we found the glutathione system may be a major regulator of protein homeostasis as observed in *C. elegans*, yeast and mammalian cell models. Our data provide a new conceptual and experimental model to address the molecular mechanisms underlying redox regulation of proteostasis. Given the evolutionary conservation of this function, we postulate that strategies aimed to maintain glutathione redox homeostasis may have a therapeutic potential in diseases associated with protein aggregation, such as most prevalent neurodegenerative diseases.

## MATERIALS AND METHODS

### *C. elegans* strains

The standard methods used for culturing and maintenance of *C. elegans* were as previously described ^79^. A list of all strains and plasmids used and generated in this study is provided in **Supplementary Table 1**. All VZ strains are 6x backcrossed with N2 wild type. All transgenes used in this study are stably expressed from genomic integrated lines, except for strains carrying the *vzEx169* and *vzEx170* extrachromosomal arrays. Details on PCR, sequencing or restriction enzyme genotyping of the different alleles as well as plasmid constructs used in this work can be provided upon request. Unless otherwise noted, all experiments were performed on synchronized worms generated by allowing 10 to 15 gravid hermaphrodites to lay eggs during two to three hours on seeded plates at 20°C.

### *S. cerevisiae* strains

A list of all yeast strains and plasmids used and generated in this study is provided in **Supplementary Table 2**. Yeast transformation was performed using a standard lithium acetate/polyethylene glycol method and all our experimentation used freshly transformed cells.

### Mammalian cell culture

Human SH-SY5Y cells were routinely grown at 37°C in a humidified incubator with 5% CO_2_ in DMEM F12+Glutamax, supplemented with 10% fetal bovine serum. All experiments were performed after 24 h of incubation. To explore whether BSO and DEM were able to induce intracellular aggregates of amyloid-beta, cells were exposed to BSO (75 mM) and DEM (25 μM) for 18 h.

### Glutathione pathway inhibitors

Appropriate volumes from stock solutions of BSO (Sigma B2515, 200 mM in H_2_O), BCNU (Sigma C0400, 20 mg/ml EtOH) and DEM (Sigma D97703, directly from the supplier, diluted in 0,5% DMSO final concentration on plates) were added to NGM prior pouring the plates. Plates were freshly prepared to avoid degradation of the chemicals with time. The developmental stages were scored in a SMZ645 Nikon stereoscope at specified days (depending on the growth temperature and genotype) after synchronized egg-lay.

### Quantification of polyQ *C. elegans* strains phenotypes

#### polyQ::YFP aggregates quantification

For worms expressing the *rmIs132 [Punc-54::Q35::yfp]* and *rmIs133 [Punc-54::Q40::yfp]* transgenes, muscle YFP aggregates were scored in the whole animal. Neuronal YFP aggregates in worms expressing the *rmIs237 [Prgef-1::AT3v1-1::Q75::yfp]* and *rmIs263 [Prgef-1::AT3v1-1::Q130::yfp]* transgenes were scored in the ventral nerve cord segment from the pharynx to the vulva. For worms expressing the *drIs20 [Pvha-6::Q44::yfp]* transgene, we scored the number of animals with 10 or more distinct YFP aggregates in the intestinal cells. All YFP aggregates quantifications were performed at 40x (muscle or intestinal aggregates) or 100x (neuronal aggregates) magnification in an Olympus BX61 fluorescence microscope equipped with a DP72 digital camera coupled to CellSens Software for image acquisition and analysis.

#### Mechanosensory response

The mechanosensory function in worms expressing the *igIs245 [Pmec-3::htt57-19Q::cfp] and igIs1 [Pmec-3::htt57-128Q::cfp]* transgenes was assayed by the touch response assay ^80^. Briefly, first day adult worms were gently touched on their tail with an eyelash mounted on a toothpick and their forward movement response was scored by 10 touches per worm assayed.

#### Time-lapse recording of blebbing and exploding embryo phenotypes

For the DIC recordings embryos were prepared and mounted as described ^81^. Gravid hermaphrodites were dissected, and two-to four-cell-stage embryos were mounted on 4% agar pads in water and sealed with Vaseline. 4D-microscopy (multifocal time lapse microscopy) was carried out using a Leica DM6000 Microscope fitted with Nomarski optics. Recordings were made using a 100X/1.4 PL APO objective, and the temperature was kept constant at 25°C. The microscope was controlled with the open-source software Micro-manager. Images from 30 focal planes (1micron/section) were taken every 30 seconds for up to 15 hours. Images and videos were processed with the Xnview and Quicktime software. For fluorescence recordings, animals were grown at 16°C and shifted to 25°C approximately 24h prior to observation. Gravid hermaphrodites were dissected and *gsr-1(m+,z+)* and *gsr-1(m-,z-)* embryos were carefully placed side by side on a ∼2% agarose pad in either H_2_O (**Figure 1g**) or M9 (**Figure 1h**). A coverslip was placed on top of the embryos and sealed with VALAP ^82^. Confocal images were acquired on a Nikon A1R microscope through a 60x/1.4 PlanApo objective using a pinhole of 1.2 Airy Units (39.7 μm); 3-5 focal planes separated by 2-3 μm were recorded every 3-10 minutes for a total of 16 h at 22-24°C. For mounting videos, a single focal plane was selected for each time point.

### Quantification of β-amyloid *C. elegans* strains phenotypes

#### aralysis

Worms expressing the *dvIs2 [Punc-54::human Aβ]* or *dvIs14 [Punc-54::human Aβ]* transgenes (and respective controls) were grown at 16°C and paralysis scoring was initiated four days after egg-lay and determined daily, whereby paralyzed worms were removed from plates. A worm was scored as paralyzed if it did not respond to a gentle touch stimulus with a platinum wire.

#### Internal hatching

Animals carrying the *dvIs14 [Punc-54::human Aβ]* transgene were grown at 16°C and the internal hatching phenotype was determined after adulthood by scoring their inability to extrude embryos, revealed by the absence of laid embryos in the area surrounding the vulva (as these animals are paralyzed). Images were captured using a SONY CCD camera (model DMK 31AU03) with the ImagingSource software (Imaging Control).

#### Developmental delay

Worms expressing the *gnaIs1 [Pmyo-2::yfp]* and *gnaIs2 [Punc-119::human Aβ; Pmyo-2::yfp]* transgenes were grown at 20°C and their size was determined three days after egg-lay. Worms were imaged at 10x magnification in an Olympus BX61 fluorescence microscope equipped with a DP72 digital camera coupled to CellSens Software for image acquisition and analysis. ImageJ Software was used to quantify the length of the worms.

### Quantification of *α*>-synuclein *C. elegans* strains phenotypes

#### *α*-synuclein::YFP aggregates quantification

For worms expressing the *rmIs126 [Punc-54::yfp]* and *pkIs2386 [Punc-54::α-synuclein::yfp]* transgenes, the YFP aggregates were scored in the head muscle cells located within the segment encompassed by the two pharyngeal bulbs.

#### Dopaminergic neurodegeneration

Worms expressing the *vtIs7 [Pdat-1::gfp]* and *baIn11 [Pdat-1::α-synuclein; Pdat-1::gfp]* transgenes were scored for neurodegeneration of the six anterior dopaminergic neurons (four CEP and two ADE dopaminergic neurons) according to previously described criteria ^36^. Briefly, worms were scored as wild type when all six DA neurons were present and their neuronal processes were intact while worms having DA cell body loss were scored as neurodegeneration.

YFP aggregates quantification and DA neuronal integrity were assessed at 40x magnification in an Olympus BX61 fluorescence microscope equipped with a DP72 digital camera coupled to CellSens Software for image acquisition and analysis.

#### Paralysis

Worms expressing the *baIn12 [Punc-54::α-synuclein::gfp]* transgene (and respective controls) were grown at 20°C and paralysis scoring was initiated four days after egg-lay and determined daily, whereby paralyzed worms were removed from plates. A worm was scored as paralyzed if it did not respond to a gentle touch stimulus with a platinum wire.

### Quantification of metastable protein *C. elegans* strains phenotypes

#### *unc-52(ts)* paralysis

Worms carrying the *unc-52(e669su250)* mutation were grown at 16°C and paralysis was scored at the specified days after egg-lay, whereby paralyzed worms were removed from plates. A worm was scored as paralyzed if it did not respond to a gentle touch stimulus with a platinum wire.

#### *let-60(ts)* embryonic arrest

Worms carrying the *let-60(ga89)* mutation were allowed to lay eggs at 16°C during 2-3 hours. Parents were removed, laid embryos counted and then allowed to grow for three days at 16°C. The embryonic arrest ratio was determined by dividing the number of developed larva by the total number of laid embryos.

### HLH-30::GFP subcellular localization

Worms expressing the *sqIs17 [Phlh-30::hlh-30::GFP]* transgene were scored at 40x magnification at the first day of adulthood for cytoplasmic versus nuclear GFP localization in an Olympus BX61 fluorescence microscope equipped with a DP72 digital camera coupled to CellSens Software for image acquisition and analysis. For the starvation assay, animals were incubated 1 hour in M9 buffer at 20°C with gently shaking prior microscopy analysis.

### SQST-1::GFP aggregates quantification

Worms expressing the *bpIs151 [Psqst-1::sqst-1::GFP]* transgene were allowed to lay eggs at 20°C for 2-3 hours. Parents were removed and embryos were incubated during 6 hours. Embryos with GFP aggregates were scored at 100x magnification in the same microscopy setting as above. For the starvation assay, pregnant adults were incubated in M9 buffer at 20°C with gently shaking during 24 hours. Embryos accumulated in the uterus of the starved animals were freed by dissecting the parents with a needle and scored as described above. For the RNAi assay, worms expressing the *bpIs151* worms were grown on empty vector control or *lgg-1* RNAi plates for 3 generations. Adults were allowed lay embryos at 20°C for 2-3 hours, further incubated during 8-10 hours until hatching and L1 larvae with more than 20 intestinal GFP aggregates were scored at 100x magnification as described above.

### *S. cerevisiae* strains phenotypes

Yeast cells were grown at 30°C by incubation in a rotary shaker using SC medium (0.67% yeast nitrogen base, 2% glucose plus drop-out mixture and auxotrophic requirements). Experiments were performed with cell cultures growing at exponential phase (OD_600_= 0.5) or diauxic phase (12h after exponential phase). To image polyQ-gfp expression and aggregates wide-field fluorescence microscopy was performed on an Olympus fluorescence DP30 BW microscope (Olympus, Tokyo, Japan) equipped with a 488 nm laser excitation for GFP. Aggregation index was determined as originally described ^83^. Cell growth was determined in a Microplate Spectrophotometer PowerWave XS (Biotek) apparatus and values were analyzed with Gen5 1.06 software. Cells (500 μl) were grown at 30°C with continuous shaking; absorbance at 600 nm was measured each 30 min for 20 hours. Resistance to disulfide stress was tested on cells grown in SC at exponential or diauxic phase. Viability was measured by plating serial dilutions (1:10) on SC plates without or with diethyl maleate (DEM) at the indicated concentration.

### SH-SY5Y cells immunofluorescence

Cells were seeded in Poly-D-lysine-coated round glass coverslips. After the treatments, they were fixed in 4% PFA for 10 minutes. Fixed cells were then blocked for 30 min in PBS (Thermo-Fisher, Massachusetts, USA) with 0.1% Triton-X and 1% bovine serum albumin. Afterwards, cells were washed three times with PBS and incubated overnight with the primary rabbit anti-Aβ42 (1:100; Lodeiro, M et al https://doi-org.proxy.kib.ki.se/10.1093/gerona/glw073). The day after, cells were washed with PBS and incubated with the secondary antibody Alexa Fluor 633 goat anti-rabbit (1:1000, Invitrogen, Corporation, Carlsbad, CA, USA) in PBS for 1h at room temperature. DAPI was used for nuclear staining. For aggregation analysis, cells were then washed thoroughly, and incubated with 0.1% (w/v) Thioflavin S in PBS for 30 min to stain for amyloid fibrils. After washing three times with PBS, cells were mounted using ProLong Gold Antifade Reagent (Thermo-Fisher, Massachusetts, USA). The primary antibody was omitted as a negative control. Confocal imaging was performed with a Zeiss LSM 510 META confocal laser scanning system. For Aβ 1-42 quantifications, Alexa Fluor 633 fluorescence was recorded with a Zeiss LSM 510 METAL confocal laser scanning system with 20x lens (NA 0.8) and 100x (NA 1.45) lens. Quantification of fluorescence intensity in individual cells was performed with ImageJ Software. For representative figures of intracellular Aβ 1-42 aggregates, Alexa Fluor 633 fluorescence and Thioflavin-S were recorded through separate channels with 100×; (NA 1.45) oil objective. Image processing was performed with the included ZEN software.

## ACKNOWLEDGEMENTS

We thank CGC (*Caenorhabiditis* Genetics Center) and NBRP (National BioResource Project), Richard Morimoto, Christian Pohl, Jan Gruber, Guy Caldwell, Randy Blakely, Hong Zhang, Enriq Herrero, Anne Bertolotti and Susan Lindquist for providing strains and plasmids. Cristina Ayuso García is acknowledged for technical assistance, Leticia Lemus for help with yeast viability assays. The Spanish Ministry of Economy and Competitiveness supported EFS and VG (BFU2016-78265-P), PA (BFU2016-79313-P and MDM-2016-0687) and AMV (BFU2015-64408-P). AMV was also supported by the Instituto de Salud Carlos III (PI11/00072). All projects were cofinanced by the Fondo Social Europeo (FEDER). AMV is a member of the GENIE and EU-ROS Cost Actions of the European Union.

## AUTHOR CONTRIBUTION

DGG, JAML and AMV designed the study, performed most experiments using *C. elegans*, analyzed the data and wrote the manuscript. BSN and JC performed the embryo recordings and quantified the blebbing/exploding phenotypes. FJNG and FML carried out the initial candidate RNAi screen and generated several strains. CPF and ACM performed the mammalian cell experiments. CDL generated some strains and performed paralysis and DEM toxicity experiments. CN contributed essential reagents and analysed the data. MDS and RPVM generated *C. elegans* strains and carried out mechanosensory assays. EFS, VG, RP and EC contributed with the yeast experiments. PA performed the embryo confocal live imaging and immunostaining. All authors edited and revised the manuscript.

## MOVIE LEGENDS

**Movie 1**

Movie recording of *C. elegans* wild type *vs gsr-1(m-,z-)* embryos. Developmental arrest of the *gsr-1(m-,z-)* embryo occurs at the pregastrula stage. Movie 1 corresponds to Fig. 1c.

**Movie 2**

Movie recording of *C. elegans rmIs133 [Punc-54::Q40::yfp]* embryos, that develop normally. Movie 2 corresponds to Fig. 1d.

**Movie 3**

Movie recording of a *C. elegans rmIs133 [Punc-54::Q40::yfp]*; *gsr-1(m-,z-)* embryo. Note the extensive cell blebbing once the embryo arrests at the pregastrula stage. Movie 3 corresponds to Fig. 1e.

**Movie 4**

Movie recording of *C. elegans rmIs133 [Punc-54::Q40::yfp]*; *gsr-1(m-,z-)* embryos. Note that the right embryo blebs and explodes while the left embryo explodes without undergoing blebbing. Movie 4 corresponds to Fig. 1f.

**Movie 5**

Movie recording of *C. elegans rmIs133 [Punc-54::Q40::yfp]*; *zbIs2 [pie-1::lifeact-Cherry]*; *gsr-1(m+,z+)* and *gsr-1(m-,z-)* embryos observed by confocal microscopy. Note that the *gsr-1(m-,z-)* embryos explode while the *gsr-1(m+,z+)* embryo completes embryogenesis and hatches. The asterisk indicates an arrested embryo that was osmotically damaged due to incomplete eggshell formation at the moment of dissection. Time is indicated in hours:minutes. Movie 5 corresponds to Fig. 1g.

**Movie 6**

Movie recording of *C. elegans rmIs133 [Punc-54::Q40::yfp]*; *ltIs37 [Ppie-1::mCherry::his-58]*; *ojIs1 [Ppie-1::GFP::tbb-2]*; *ltIs24 [Ppie-1::GFP::tba-2]*; *gsr-1(m+,z+)* and *gsr-1(m-,z-)* embryos observed by confocal microscopy. The left part displays GFP::tubulin whereas the middle part corresponds to mCherry::histone H2B. Abnormal accumulation of microtubules at the plasma membrane and perinuclear chromatin condensation is particularly evident in the *gsr-1(m-,z-)* embryo at the top. Time is indicated in hours:minutes. Movie 6 was acquired under similar conditions as the still images represented in Fig. 1h.

**Movie 7**

Movie recording of *C. elegans* wild type *vs ced-10(t1875)* embryos. The developmental arrest of the *ced-10(t1875)* embryo occurs at the premorphogenesis stage, while *gsr-1(m-,z-)* embryos arrest earlier, at the pregastrula stage (Movie 1). Note that the blebs formed by *ced-10(t1875)* embryos are not as large and prominent as those observed in *gsr-1(m-,z-); rmIs133* embryos (Movies 3-4).

